# Intravital microscopy and statistical image analysis of early engraftment of haematopoietic stem cells reveal dynamic and evolving cellular behaviours linked to progressive differentiation

**DOI:** 10.64898/2026.09.11.750961

**Authors:** Sara Gonzalez-Anton, Chang Liu, Reema Khorshed, Mari Carmen Romero-Mulero, George Adams, Ben Partridge, Alvin Ziqi Lu, Nina Cabezas-Wallscheid, Constandina Pospori, Chiu Fan Lee, Ken R. Duffy, Cristina Lo Celso

**Author notes:** Corresponding author: Cristina Lo Celso. authors contributed equally to this work (Sara Gonzalez-Anton, Chang Liu).

## Abstract

Haematopoietic stem cells (HSCs) have long been used in the clinic for bone marrow transplantation applications, critical for the survival of an increasingly wide range of patients with haematological, oncological and immunological pathologies. Despite this, little is known about the mechanisms through which relatively few stem cells are able to regenerate the entire haematopoietic tissue of transplant recipients. To gain insights on this biological process, we used intravital microscopy of calvarium bone marrow, collecting tissue-wide images and a total of 850 hours of tracks of engrafting haematopoietic stem and progenitor cells from 24 hours to 8 days following injection in lethally irradiated recipients. Analysis of the data revealed that regenerating HSCs and their immediate progeny are highly dynamic, migrating through the parenchyma at both the microscopic and near-macroscopic scales, i.e. within and in-and-out of fields of view. HSC-derived cell clusters, expanded locally between day 2 and day 4 post-transplant, leading to patches of densely populated bone marrow by day 8. Single cell level analysis highlighted increasingly heterogenous cellular behaviours over time. Track clustering based on migration and niche interaction parameters combined with post-tracking whole mount immunostaining revealed that persistence in the vicinity of nestin-GFP perivascular cells and relatively slow movement correlated with stemness, while more heterogeneous niche interactions coupled with slowest or faster migration correlated with differentiation, validated through flow cytometry analysis and functional studies. The findings presented here shed light on the fundamental principles driving haematopoietic regeneration in transplantation settings.

## INTRODUCTION

Haematopoietic stem cells (HSCs) have an essential role in the adult life of vertebrates, sustaining constant blood production throughout life. HSCs are defined by their multipotency and self-renewal capacities (Till and McCulloch, 1961). With their ability to give rise to all other blood lineages, while also keeping a pool of themselves, HSCs were the first stem cells to be exploited in the clinic, even prior to the identification of specific cell surface markers that allowed their purification (Mathé et al., 1959; Thomas et al., 1957). HSC transplantation (HSCT) is used as treatment for many diseases, mainly haematological malignancies, but also some other haematological disorders, solid cancers, and an increasing number of autoimmune diseases (Kanate et al., 2020). Despite the wide application of HSCT as treatment, we still do not completely understand this complex process. While engraftment is most often achieved, all recipients live through a few weeks of cytopenia. 5-10% of transplant recipients fail to reconstitute one or two blood lineages, and for those few with complete engraftment failure the mortality rate can be as high as 80% (Sun et al., 2021). A better understanding of the biological processes underpinning bone marrow (BM) regeneration following HSCT, and especially of those enabling HSCs to drive it, is needed to increase the rate of multilineage recovery, reduce the duration of cytopenia, and improve transplantation outcome.

The haematopoietic cell hierarchy has been extensively studied at the molecular and functional levels up to single cell resolution. At the top of the pyramid are HSCs and multipotent progenitors (MPPs), giving rise to increasingly proliferative and committed progenitors, eventually replenishing all the terminally differentiated cells (Laurenti and Göttgens, 2018). Several markers have been identified to phenotypically describe and characterise primitive stem and progenitor cell populations (HSPCs) and their progeny (Kiel et al., 2005; Pietras et al., 2015), and over the last decades several studies have started to analyse the contribution of different progenitor subpopulations to haematopoiesis, in both homeostasis and stress. It has been shown that MPPs have an important role contributing to blood cell numbers during steady state (Busch et al., 2015; Sun et al., 2014), however HSCs are the only population able to achieve long term reconstitution following transplantation (Dykstra et al., 2007; Sawai et al., 2016). It is widely accepted that HSPC function is the result of cell intrinsic and extrinsic regulatory factors. Population-level and single cell transcriptomic analysis have provided a detailed understanding of cell intrinsic mechanisms underpinning HSPC function, and microscopy studies have been pivotal in tackling the complexity of cell-extrinsic mechanisms.

Several bone marrow microenvironment components, including mesenchymal stromal cells, endothelial cells, immune cells and neuronal terminations, have been demonstrated to be key for HSC maintenance and function, and are therefore considered to be part of the HSC niche (Comazzetto et al., 2021; Pereira et al., 2024). Several questions remain open about the nature of HSC-niche interactions, especially in the context of transplantation. In particular, we need to understand not only what cell types interact with HSCs, but the very nature of these interactions, i.e., whether they are stable or transient, and in the latter case how long lasting they may be. Understanding these details is critical to enable the development of therapeutic interventions to regulate HSC output.

Intravital microscopy (IVM) provides the unique opportunity to follow cellular and tissue changes longitudinally within the same mouse during time periods ranging from hours to weeks. IVM of murine bone marrow has been an essential tool to uncover the nature of HSPC-niche interactions, including during haematopoietic stress and transplantation settings (Adams et al., 2009; Christodoulou et al., 2020; Lo Celso et al., 2009; Rashidi et al., 2014; Sanchez-Aguilera et al., 2011; Secchi et al., 2025; Upadhaya et al., 2020). These studies have highlighted how HSPC-niche interactions are both heterogenous and temporally dynamic and opened several further questions, such as whether there may be spatiotemporal cell-niche interaction patterns specific to HSCs and their progeny, whether cell-niche interactions may change gradually as differentiation occurs, similarly to what has been described for the transcriptome of early HSPCs. To address these questions in the context of HSCT, here we performed IVM on Nestin-GFP stroma reporter mice and Flk1-GFP endothelial reporter mice that had been lethally irradiated and injected with purified HSC and MPP cell populations. By following the cellular dynamics developing over up to 8 days from injection, we were able to analyse the early stages of HSPC engraftment and haematopoietic regeneration. As a result, we were able to cluster cells based on their migratory and niche interaction characteristics, to identify patterns specific to HSCs and their progeny and to highlight how progressive differentiation is linked to gradual changes in cell-niche interactions. Our findings allow the identification of HSPC populations in IVM datasets based on their behaviour rather than a multitude of cell surface markers or transcriptomics-based scores and improve our understanding of the cellular dynamics leading to haematopoietic regeneration following HSCT.

## MATERIALS AND METHODS

### Mice

All animal work was performed in accordance with the UK Home Office regulations under the Animals (Scientific Procedures Act) 1986 and the Animal Welfare and Ethical Review Body (AWERB) guidelines at Imperial College London and Sir Francis Crick Institute. All procedures were approved by the UK Home Office under project license PP9504145.

All mice were bred and housed at Imperial College London or Sir Francis Crick Institute, according to institutional guidelines. mT/mG mice (Muzumdar et al., 2007) were obtained from Sir Francis Crick Institute. Flk-1-GFP mice (Xu et al., 2010) were a gift from Alexander Medvinsky (University of Edinburgh); Nestin-GFP mice from Paul Frenette (Einstein School of Medicine, New York) (Mignone et al., 2004); Flk2-cre mice were a gift from Camilla Forsberg (University of California, Santa Cruz) (Benz et al., 2008; Boyer et al., 2011); Col2.3-CFP mice were a gift from David W. Rowe (University of Connecticut Health Center) (Hawkins et al., 2016; Kalajzic et al., 2002; Paic et al., 2009). For all experiments and transplantation assays, cohorts of male mice between 11 to 16 weeks were used.

### HSC and MPP purification and transplantation

All transplantation experiments were carried out under strict sterile conditions. Bones from healthy mT/mG or mT/mG x Flk2-cre donor mice were harvested (femurs, tibias, iliac bones, sternum, and vertebrae), gently crushed in FACS buffer (2% Foetal Bovine Serum (FBS; Life Technologies, Cat N° 10500-064) in Dulbecco’s Phosphate Buffered Saline, without calcium chloride and magnesium chloride (PBS; Sigma Aldrich, Cat N°D8537)) using a mortar and a pestle, pipetted into a single cell suspension, and filtered through a 40 μm strainer. The samples were then centrifuged at 500g for 5 minutes at 4°C, resuspended and incubated for 5 minutes in red blood cell lysis buffer (0.001g/ml Potassium bicarbonate (Cat N° 60339), 0.00 8g/ml Ammonium chloride (Cat N° A9434), 20mM EDTA (Cat N° EDS) (all from Sigma), and 5% (v/v) FBS in Milli-Q water) at room temperature. After that time, samples were centrifuged again at 500 g for 5 minutes at 4°C. The resulting single cell suspensions were stained with the biotinylated lineage cocktail (CD3, CD4, CD8, Ter119, B220, Gr-1 and CD11b). Samples were lineage depleted using streptavidin magnetic microbeads (130-048-101; Miltenyi Biotech), LD columns (130-042-901; Miltenyi Biotech) and magnets (Miltenyi Biotech). The lineage depleted samples were then centrifuged and stained with the following antibody panel: Fixable Viability 780, Streptavidin BV510, c-Kit PerCP Cy5.5, Sca1 BV711, CD48 PE Cy7, CD150 BV650, CD34 APC, Flk2 CF-594. Single colour controls for compensation were prepared using OneComp eBeads (01-1111-42; Biolegend) or WT BM cells. Fluorescence-minus-one (FMOs) controls were prepared when necessary, using sample cells and following same staining protocol.

All samples were sort purified using a FACSAria III (BD Biosciences). The sorted cells were injected intravenously in recipient mice that had been conditioned with lethal irradiation administered in two doses of 5.5 Gy 3 hours apart earlier that day. Sort purified cells (7,000 to 21,000 HSCs and 140,000 or 200,000 MPPs) were transplanted intravenously into each mouse, 3-4 hours after irradiation.

### Intravital microscopy

Surgery and BM calvarium intravital imaging were performed as previously described (Duarte et al., 2018b; Haltalli et al., 2020; Hawkins et al., 2016; Lo Celso et al., 2009; Pirillo et al., 2022). Mice were anaesthetised using isoflurane in medical oxygen, using 4% isoflurane in 2 L/min O2 for induction and 2-1% isoflurane and 2L/min O2 for maintenance. Minimal invasive surgery was performed to remove the skin on top of the skull (calvarium) and expose the skull bone. Using Diamond Carve Dental Cement (Associated Dental Products; Cat. N° SUN527) a small custom-made metal headpiece (imaging window) was attached to the bone. Once the cement was completely solidified, the mouse was transferred to the microscope stage. The mouse was placed on a heated pad, and its temperature was constantly monitored through a thermal probe connected to a thermal sensor. To protect the eyes from dryness and corneal injury a lubricant eye ointment was applied (Lacri-lube®). The lock and key mechanism linking the headpiece to the mouse holder positioned onto the stage, allowed continuous stable imaging of the calvarium BM. A small amount of PBS was placed on the window to soften the subcutaneous tissue on top of the bone, which was then cleaned with a cotton bud. PBS was added for imaging acquisition. Blood vessels were visualised by injecting 80 µl of 8 mg/ml Cy5-Dextran (500KDa; NANOCS, DX500-S5-1) intravenously.

IVM was performed using an LSM 780 Zeiss upright confocal/two photon hybrid microscope equipped with Argon laser (458, 488 and 514 nm), a diode-pumped solid-state 561 nm laser, a Helium-Neon laser (633 nm), and tuneable infrared multiphoton laser (Spectraphysics Mai Tai DeepSee 690-1020 nm), 4 non-descanned detectors (NDD) and an internal spectral detector array. The signal was visualised using a Zeiss W Plan-Apochromat x20 DIC water immersion lens (1.0 N.A.). An external dichroic mirror (450nm) was used when confocal and two photon microscopy modalities were combined.

Second harmonic signal was used to visualise the bone and was excited with the two-photon laser tuned at 870nm and detected with external detectors. GFP signal was excited at 488nm, tdTomato at 561nm and Cy5 was excited with 639nm, and all of them were detected using the internal detectors.

Acquisition of 3D tile scans of the whole BM area within the imaging window, defined by coronal suture, central sinus, bifurcation, and lateral bone was achieved by stitching of adjacent z-stack images. The tile scans allowed the visualisation and analysis of calvarium BM at single cell resolution. Acquiring specific positions of interest within the tile scan repeatedly over time (3 minutes interval between timeframes) allowed to produce time lapse datasets to study cell dynamics and interactions.

When it was necessary to image the same fields of view during different imaging sessions, mice were subcutaneously injected with analgesic (Buprenorphine 0.1 mg/kg) one hour prior to the end of the first imaging session. The imaging window was covered with intrasite gel (Intrasite gel applipak (8 g), Smith&Nephew, Cat N° 66027308) to protect the bone and prevent scar formation, and bandaged. Mice were allowed to recover from anaesthesia and placed in a separate cage, where buprenorphine mixed in raspberry jelly (0.8 mg/kg) was provided. Mice could then be re-anaesthetized, placed on the microscope stage and the same positions could be found and re-imaged.

### Flow cytometry

BM samples were obtained in the same way as for HSC and MPP purification and stained for 20 minutes at 4 °C in the dark with a lineage antibody cocktail (1:40), which includes the following biotinylated antibodies: CD3, CD4, CD8, Ter119, B220, Gr-1 and sometimes CD11b, mixed at a 1:1 ratio. After staining, the cells were washed in FACS buffer and centrifuged at 500 g for 5 minutes. The sample was then resuspended in an antibody mix prepared in FACS buffer, containing all fluorophore-conjugated antibodies (supplementary table 1) and streptavidin (1:1000). All conjugated antibodies were diluted 1:200 apart from CD34 (1:80) and Flk2 (1:100). After staining, the cells were washed in FACS buffer, centrifuged at 500 g for 5 minutes at 4 °C and resuspended in FACS buffer for acquisition. 4,6-diamidino-2-phenylindole (DAPI) (NucBlue® Fixed Cell ReadyProbes® Reagent; Invitrogen, Cat. # R37606) was used to assess viability and separate live and dead cells.

Single colour and FMOs controls were prepared as for HSC and MPP purification.

For the analysis of peripheral blood reconstitution, 50-100 μl of peripheral blood were collected by venipuncture into EDTA coated tubes. Red blood cells were lysed as described above, and cells were stained with the following antibodies: CD45, CD11b, CD3 and B220. Recipients were considered reconstituted if >0.5% peripheral blood cells were donor-derived at 20 weeks post-transplant.

All the samples were acquired with a LSR Fortessa (BD Biosciences), and the data were analysed with FlowJo (Tree Star) software.

### Wholemount staining

Calvaria were collected from mice after IVM sessions and fixed overnight in 4% PFA. Samples were always kept at 4 °C, incubated in a blocking buffer (5% goat serum) for 24 hours, then with primary antibodies (biotin-conjugated lineage antibody cocktail same as used for FACS, diluted 1:50; biotin-conjugated anti-CD41, 1:100; biotin-or FITC anti-CD48, 1:100; AF647-conjugated anti-CD150, 1:50) for 48 hours, washed in PBS overnight, incubated with AF405-conjugated streptavidin (1:200) for 3-5 days and washed overnight again. Stained samples were kept in PBS for microscopy analyses, which were performed using the same settings as for intravital microscopy.

### Image processing and cell tracking

ZEN Black (Zeiss, Germany) software was used to stitch three-dimensional BM tile scans.

Time lapse datasets were 4D hyperstacks (x,y,z,and time). Before tracking, tomato signal bleaching was corrected using the Fiji bleach correction plugin. Cell tracking was performed manually using IMARIS (Bitplane, Switzerland). To increase accuracy in cell tracking data, displacements in the Z plane caused by movement artifacts were corrected by applying 4D data registration protocols implemented in FIJI (Preibisch et al., 2010) that allowed to register the time-lapse datasets before cell tracking. Coordinates from each cell at each time point were exported from IMARIS for further analysis. Number of cells per stack was counted on the first time frame image.

For figure building purposes, image processing was performed in Fiji. 3D frames were flattened into maximum projections; HSC Day2, HSC Day4 and MPP Day 2 cells were segmented by manual thresholding in both time-lapse and whole mount images, and resulting cropped signal is shown. Note nestin-GFP signal in videos containing MPP Day2 cells underwent the same process.

### Analysis of cell tracks

All tracking data were analysed using R (version 4.4.1). The following parameters were calculated for each track: mean speed, step speed standard deviation, arrest coefficient (Khan et al., 2018), maximum radius, and distance from initial to final position, overall distance covered (path length), confinement index radio (Shakhar et al., 2005) and nestin arrest coefficient (NAC). These parameters were calculated as follows.

Let *B* = {*A*_1_, *A*_2_, ⋯, *A_n_*} be the set which included *n* unique cells, where A_k_ meant the *k*-th unique cell. For 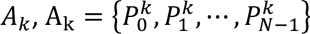, where 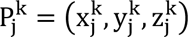 was the three-dimensional geographic coordinate for the *j*-th track record, and *N* was the number of records in the *k*-th cell. The formula for the distance in 3 mins between each record was

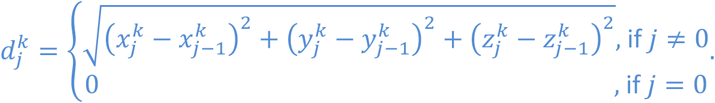

Let 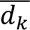 be the average speed (µ*m*/*min*), and σ*_k_* be the standard deviation of speed of the *k*-th cell

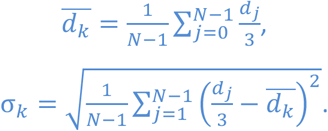

The arrest coefficient was the proportion of time the cell’s instantaneous velocity was less than 2 *μm*/*min* (Khan et al, 2018). The formula for *k*-th cell’s arrest coefficient (*a_k_*) was

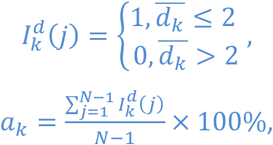

where, 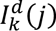 was an indicator function. Let *m_k_* present the maximum radius, and *s_k_* be the path length of the *k*-th cell movement.

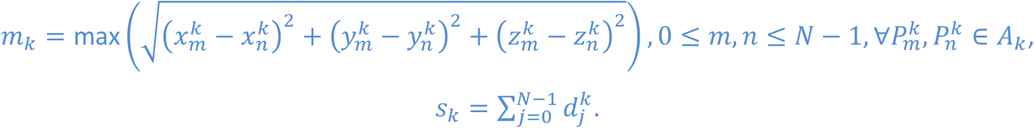

The confinement ratio and confinement index described the linearity of cell’s movement. The confinement index was calculated as the ratio of maximum radius to path length (Shakhar et al., 2005). The formula for the *k*-th cell’s confinement index (*i_k_*) was

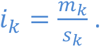

The confinement ratio (*r_k_*) was the ratio of the distance between the initial and the final positions of each cell to the total distance covered by the same cell (Hugues et al., 2004). The formula for the *k*-th cell’s confinement ratio was

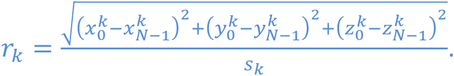

For the *k*-th cell, the shortest distance from the cell to the nestin at each 3 mins was recorded in the set 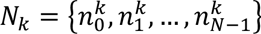, where 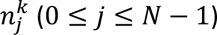 was the shortest distance from the *k*-th cell to nestin at time *j*. Nestin arrest coefficient (NAC) is the proportion of time a cell is less than 25µ*m* away from nestin. The formula for the NAC was

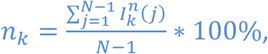

where, 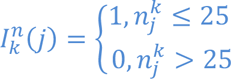 was an indictor function.

Tracks were clustered using Uniform Manifold Approximation and Projection for Dimension Reduction (UMAP) with the UMAP package in R. In order to classify cells into different clusters, mean speed, standard deviation of speed, arrest coefficient, confinement index and nestin arrest coefficient were used. Visualisations were completed using the UMAP (version 0.2.10.0) and ggplot2 (version 3.5.0) packages in R.

The Pearson correlation coefficient between tracks’ displacement and maximum radius was calculated by the ‘cor’ function with the method ‘pearson’ in R.

### Gene Ontology analysis

GO enrichment analysis of previously published HSC or MPP2/3/4 signatures (Sommerkamp et al., 2021) was performed using the clusterProfiler package with the org.Mm.eg.db annotation database. Over-representation analysis of Biological Process (BP) terms was conducted with enrichGO (p-value cutoff = 0.1). Redundant terms were collapsed with simplify (similarity cutoff = 0.7, retaining the term with the lowest adjusted p-value). Selected enriched terms were visualized with the dotplot function.

### Statistical analysis

Statistical analyses were completed using R packages. Permutation tests with Bonferroni corrections were applied to all track analyses. The two-tailed p-values were calculated by Monte-Carlo approximation with 250,000 sampling. The outcomes were treated as significant if p < 0.0167 (p < 0.0167*, p<0.0033**, p<0.00033***).

## RESULTS

### HSC-driven regeneration is initiated by localised growth of cell clusters

To study the early stages of HSC engraftment following transplantation, we used a well-established experimental model of HSCT. HSCs (Lin-c-Kit+ Sca-1+ CD48-CD150+ Flk2-CD34-) expressing tomato fluorescent protein on their membrane were sort purified from mT/mG (Muzumdar et al., 2007) transgenic mice and injected into recipients that had been conditioned by irradiation (Figure 1A, B). Recipient mice were either wild type, Nestin-GFP, Flk1-GFP or Col2.3-CFP transgenic mice to allow simultaneous observation of injected cells and either nestin-GFP positive periarteriolar stroma (Méndez-Ferrer et al., 2010), endothelial cells (ECs) (Duarte et al., 2018a), or osteoblasts (Hawkins et al., 2016; Kalajzic et al., 2002; Paic et al., 2009), respectively. These two reporter lines were selected to build on previous studies reporting engrafting HSPCs localising near osteoblasts and vasculature immediately after injection and up to two days later (Khorshed et al., 2015; Lo Celso et al., 2009; Rashidi et al., 2014). Nestin-GFP stroma cells had been reported to survive following irradiation and static IVM images had captured engrafting HSPCs in the vicinity of these cells (Méndez-Ferrer et al., 2010), however the kinetics of HSC-EC and HSC-nestin-GFP cells interactions had not been reported before. Recipient mice were analysed using hybrid confocal and two photon intravital microscopy at 1, 2, 4 and 8 days following transplantation, and tomato+ cells found in the calvarium BM cavities located between the coronal suture and central sinus bifurcation were recorded for a number of hours using time-lapse imaging (Figure 1B, C).

**Figure 1.**
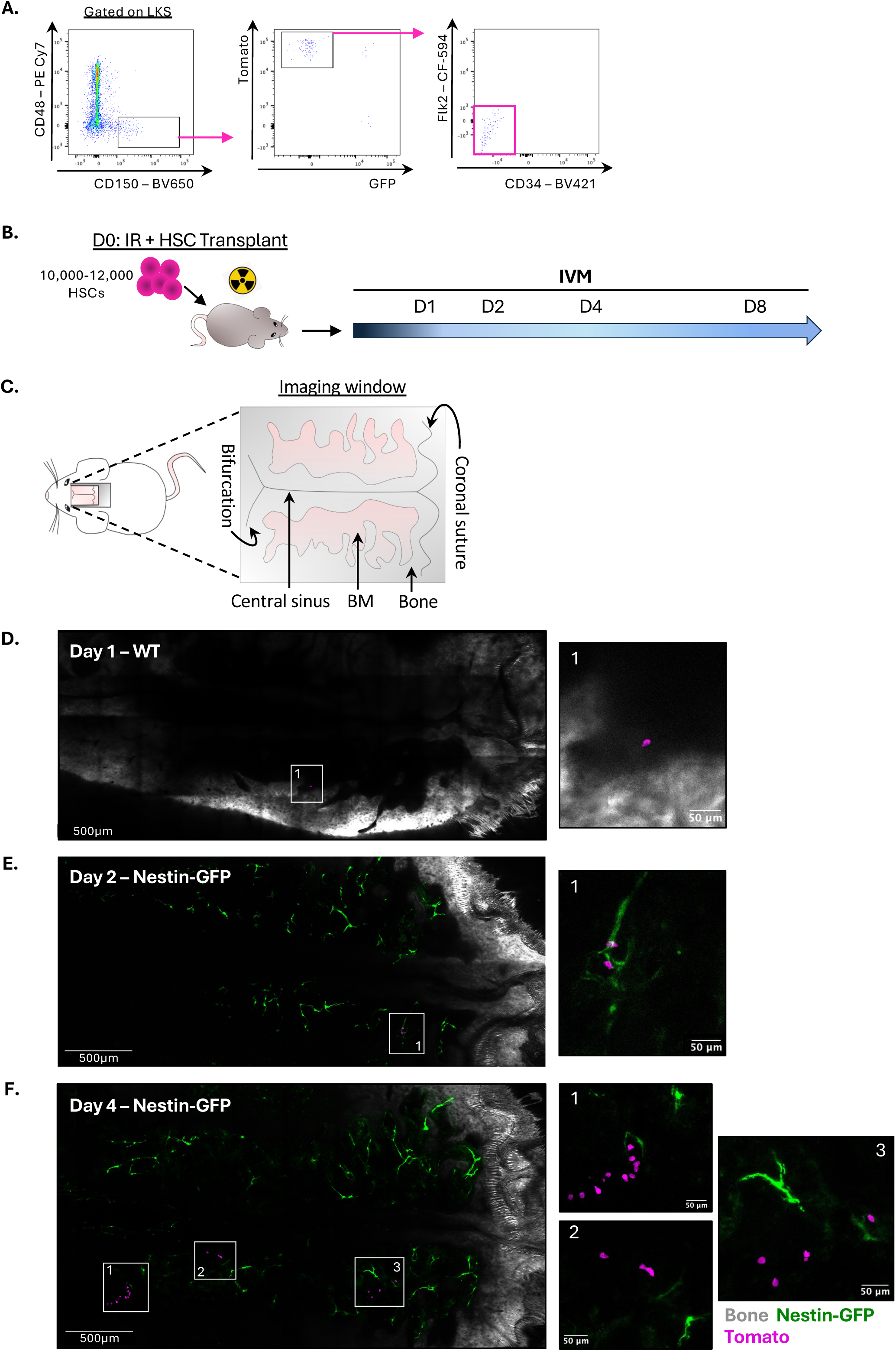
HSC-driven early engraftment is achieved through local growth of cell clusters. **(A)** Representative example of flow cytometry gating strategy used to sort-purify HSCs (gated on live, single cells, Lineage negative, c-Kit+ and Sca-1+). **(B)** Experimental timeline for HSC sort, transplant and IVM. **(C)** Schematic representation of imaging window for IVM and relevant calvarium structures. **(D, E, F)** Representative tile scan images of calvarium BM of WT/nestin-GFP mice transplanted with HSCs and imaged at day 1 **(D)**, day 2 **(E)** and day 4 **(F)** after transplantation. White framed areas within the tile scan images are shown at higher magnification on the right side to highlight tomato+ cells. White: bone collagen second harmonic generation signal; green: GFP; magenta: tomato.

When mice were imaged one day after transplantation, they typically presented a single tomato+ cell in a peri-endosteal location (Figure 1D), consistent with previous observations (Khorshed et al., 2015; Lo Celso et al., 2009; Rashidi et al., 2014). On day 2 after transplantation, we could find 1 to 6 tomato+ cells per calvarium, mostly as single cells but occasionally with 2 or even 3 cells/field of view (284μm^2^) (Figure1E). At day 4 post HSCT we could identify multiple small clusters of 4-15 clearly identifiable cells, with cells in each cluster in the vicinity of each other but only occasionally in direct contact (Figure1F). Some mice were imaged at both day 2 and day 4, and in this case fields of view with cells on day 2 contained more cells, arranged in clusters, on day 4, with further, new cell clusters identifiable in other fields of view at day 4 only. When the recipient mouse was Nestin-GFP transgenic, nestin-GFP perivascular cells were normally present within the same field of view of each cluster, at varying distances from the tomato+ cells (Figure1 E, F). By day 8 post HSCT, the clusters were much bigger, with densely packed cells that would have been too challenging to track in time lapse microscopy datasets (Supplementary figure 1A). These images indicated that HSC engraftment is driven by the homing and lodging of individual HSCs, followed by the generation of lose clusters of cells, which eventually grow into wider, densely packed cell clusters. Within this process, the first four days following injection are the ideal time-window to investigate the dynamic nature of the interactions between single HSCs, their progeny and the BM microenvironment.

We previously demonstrated that cell clusters observed during the first few days post HSCT are the result of HSC proliferation (Lo Celso et al., 2009), and it is widely accepted that HSC-driven regeneration of haematopoietic tissue is driven by fast proliferation resulting in some self-renewal but also a significant amount of differentiation (Dong et al., 2020). We therefore reasoned that the cells observed at day 2 and day 4 post HSCT had to be a mixture of HSCs and their progeny. This raised the question whether it would be possible to identify cellular behaviours specific to stem cells and to progenitor cells, and whether based on these, one could recognise which of the observed cells were stem cells and which ones were progenitor cells. To aid with this identification, in some experiments we took advantage of the cre-driven reporter switch of the mT/mG allele, generated mT/mG x Flk2-Cre double transgenic donors and sort purified GFP+ MPP4 cells (Lin-c-Kit+ Sca-1+ CD48+ CD150-Flk2+ CD34+, referred as MPP from now on; Supplementary Figure 1B) in addition to tomato+ HSCs (Figure 1A). In these experiments, both cell populations were injected in each recipient, and we tracked simultaneously HSC and MPP immediate progeny (Supplementary Figure 1C). To maximise the chance of IVM of MPP progeny highlighting cellular behaviours typical of immediate HSC progeny, we limited imaging of MPPs to day 2 following injection, i.e. at a time when HSCs have been described to generate MPPs (Dong et al., 2020). Tilescan images of recipient mice at day 2 post HSC + MPP injection revealed that many more GFP+ than tomato+ cells could be detected, with MPPs generating numerous clusters of clearly identifiable GFP+ cells (Supplementary Figure 1D). Taken together, these data indicated that IVM allows to identify and track HSC and MPP progeny during the first days following transplantation, and it is therefore the ideal tool to uncover the cellular behaviours that may be linked to HSC engraftment and differentiation.

### Post-transplant HSC progeny – niche interactions are dynamic and increasingly heterogeneous over time

To uncover the spatiotemporal nature of the interactions between HSPCs and the bone marrow microenvironment during early HSC-driven haematopoietic regeneration, we examined the 3D time lapse datasets associated with each tile scan. The time lapse datasets were generated by recording a 3D stack centred on every detected cell or cell cluster every 3 minutes for up to 11 hours (Figure 2A-F). All stacks had the same xy dimensions, corresponding to a single field of view (283µm^2^), and contained 12-14 Z slices acquired at 3-5 µm interval. In total, we recorded time lapse datasets from 13 different transplantation experiments, collecting approximately 280 hours of time-lapse images, each spanning from 1 to 11 hours in length and capturing 1 to 20 cells (Supplementary figure 2A-B). For simplicity, we will refer to HSC progeny observed at day 2 post-transplant as “HSC Day2 cells”, those observed at day 4 as “HSC Day4 cells”, and to MPP progeny as “MPP Day2 cells”. These are the three groups of cells we analysed and compared. Consistent with the analysis of the tile scans, we observed the lowest number of cells per 3D field of view for HSC Day2, and higher numbers for HSC day4 and MPP Day2 cells (Supplementary figure 2B). When both HSCs and MPPs were injected in the same recipient, some fields of view would contain both HSC and MPP progeny (Supplementary figure 2C), and others only MPP or only HSC progeny. Moreover, we observed a positive correlation between the number of cell-containing fields of view and the number of cells found per mouse (Supplementary figure 2D), indicating that HSC-driven haematopoietic reconstitution happens through focal growth in an increasing number of BM areas.

**Figure 2.**
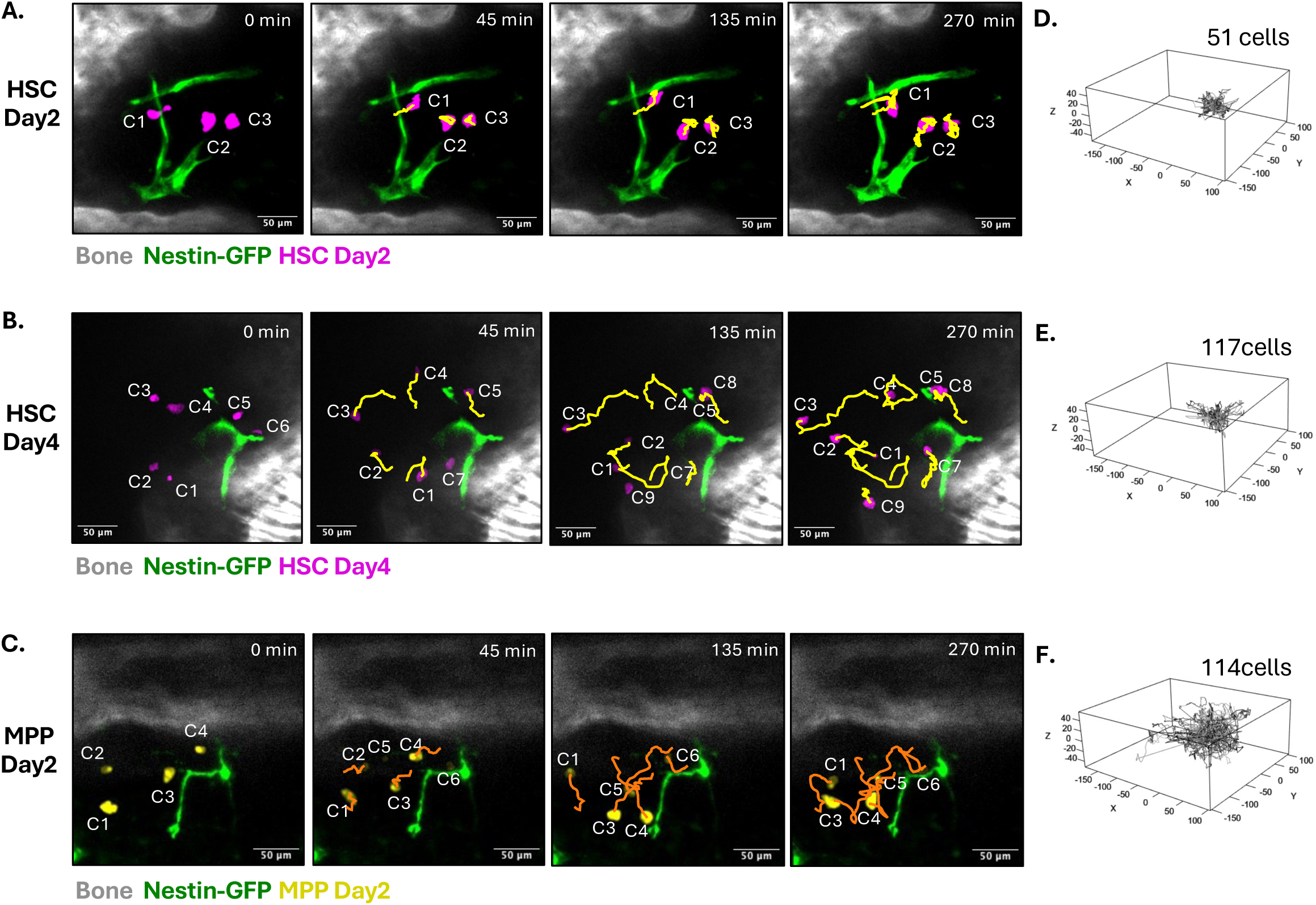
Time-lapse of HSC and MPP progeny reveals heterogeneous migratory behaviours. (A-C) Representative time frames of time lapse datasets acquired on nestin-GFP mouse calvaria, showing tracks of transplanted tomato+ HSC progeny cells at day 2 after transplant (HSC Day2) **(A)**, tomato+ HSC progeny cells day 4 after transplant (HSC Day4) **(B)** and GFP+ MPP progeny at day 2 after transplant (MPP Day2) **(C)**. **(D-F)** Rose plots showing all cell tracks of HSC Day2 **(D)**, HSC D4 **(E)** and MPP Day2 **(F)** cells.

An initial qualitative analysis of the resulting time lapse movies provided an overview of the heterogeneity of cellular behaviours detectable when observing HSC and MPP progeny at day 2 and 4 post injection (Figure 2A-C). Virtually no cell observed was completely still, and we could detect a range of behaviours, from cells that presented minimal displacement but continuously sensed the environment through active, dynamic protrusions (Figure 2A cell 3 and supplementary video 1), to localised wiggling with virtually no displacement (Figure 2A cell 2 and supplementary video 1), to varying degrees of migration through the parenchyma (Figure 2A cell 1, Figure 2B cells 1 to 9 and supplementary videos 1 and 2), and even intravasation and disappearance from the field of view (Supplementary video 3). A number of cells appeared in the field of view at varying times from the start of the time lapse. This last behaviour could result from either extravasation into the parenchyma or migration from deeper areas, and was particularly frequent for MPP Day2 cells (Supplementary video 4), as highlighted in a swimmer plot where each cell track is represented as a line that starts at the time of first appearance of the cell within the imaging session (Supplementary figure 2A). In summary, initial visual evaluation of all datasets indicated that MPP Day2 cells displayed the most movement, and HSC Day2 the least.

Next, each cell was tracked in 3D using a semi-automated tracking method based on the Imaris cell tracking algorithm, and the 3D coordinates of the position of each cell in each time frame were recorded. 282 cells were tracked in total, amounting to a total of 17237 3D positions recorded or 848 hours of tracking data. Rose plot visualisation of all recorded tracks provided a first confirmation that HSC Day2 cells were the least motile, MPP Day2 cells the most motile, and that HSC Day4 cells had an intermediate migratory phenotype (Figure 2D-F). This raised the question whether other characteristics of the tracks generated by the three groups of cells would rank in the same way.

### HSPC movement characteristics evolve gradually as differentiation occurs

To determine potential specific cell behaviours linked to differentiation state, we measured and compared multiple track parameters for each group of cells. Because not all tracks had the same duration due to differences in the length of IVM sessions, we focussed on parameters that would not be affected by the duration of the track: speed, arrest coefficient, confinement index and confinement ratio (Figure 3A-E). HSC Day2 cells had the lowest speed, and MPP Day2 cells were the fastest. Interestingly, HSC Day4 cells presented an intermediate mean speed when looking at the overall population, and had a wider distribution of values across the population compared to HSC Day2 cells (Figure 3A). The same pattern was observed for the standard deviation of speed. HSC Day2 cells had the lowest median for this parameter, and the distribution of values for this cell group was more homogeneous compared to that of HSC Day4 and MPP Day2 cells, suggesting that the HSC Day2 cell population is relatively homogeneous in terms of its movement. HSC Day4 and MPP Day2 cells had higher standard deviation of speed, indicating higher heterogeneity of cell movement within these cell populations (Figure 3B).

**Figure 3.**
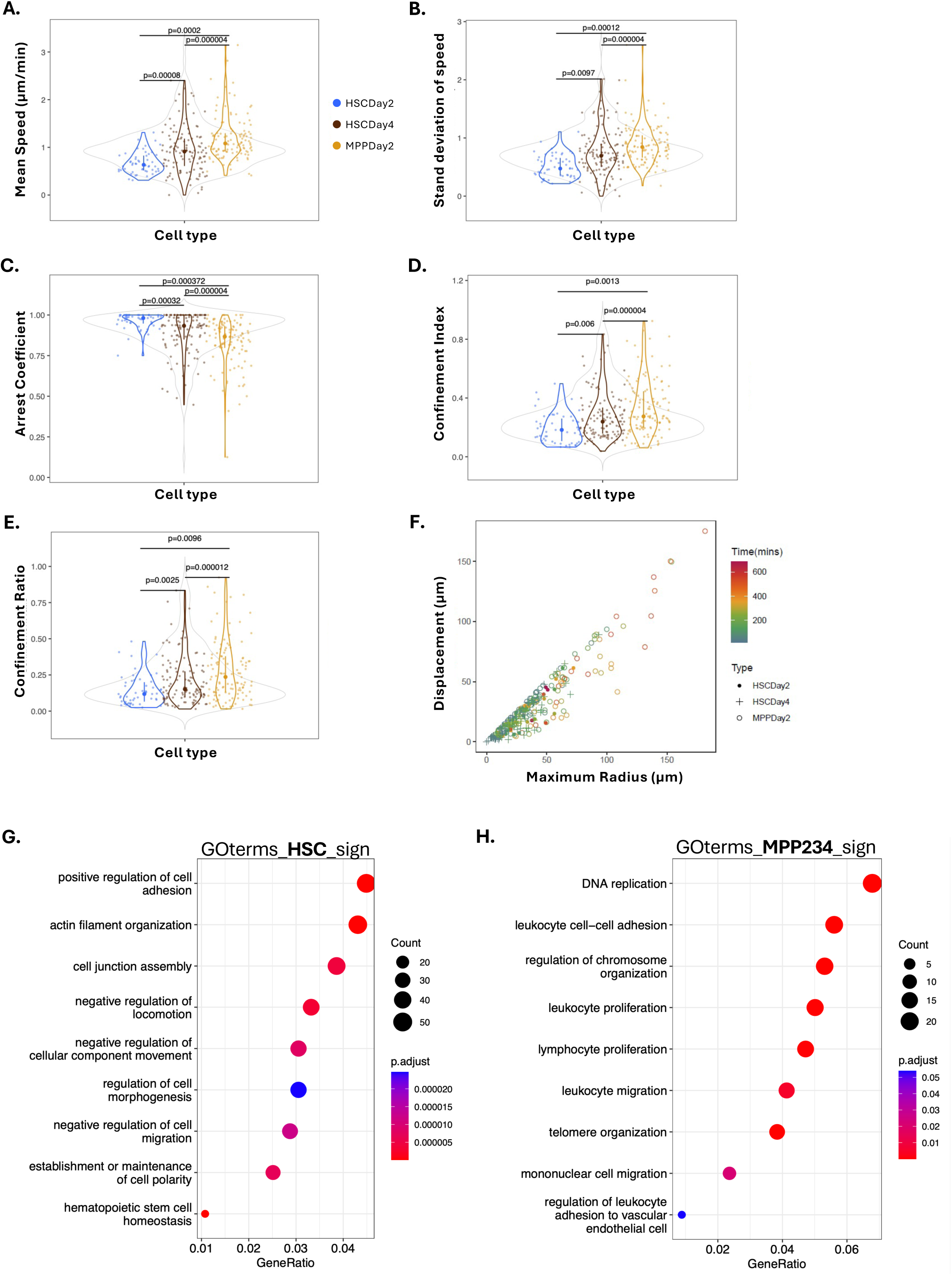
Evolving migratory behaviour from Day 2 to Day 4 and HSC to MPP progeny, consistent with transcriptional changes. (A-E) Violin and dot plots of speed (A), standard deviation of the speed (B), arrest coefficient (C), confinement index (D) and confinement ratio **(E)** of HSC Day2, HSC Day4 and MPP Day2 cells. Each dot is a tracked cell, vertical lines highlight values within the two middle quartiles for each population, and larger dots highlight median values. Grey violin plots in the background encompass the three populations combined. P values were calculated using permutation tests with Bonferroni correction. **(F)** Dot plot of displacement vs. maximum radius measurements of all tracked cells, colour coded based on track duration. **(G, H)** Gene Ontology analysis of HSC **(G)** and combined MPP2-3-4 **(H)** populations from Cabezas-Wallscheid *et al*., 2014.

The arrest coefficient is the proportion of time a cell spends without changing position. HSC Day2 cells had the highest arrest coefficient, with many cells being largely immotile (Figure 3C). These were the cells we initially identified as actively sensing the environment through dynamic protrusions rather than active migration, and those that wiggled within a highly confined space (Figure 2A, e.g. cell C3 and C2). HSC Day4 cells had a lower median arrest coefficient and a proportion of cells within this population retained a very high arrest coefficient, while others were more actively migratory and presented lower arrest coefficients. MPP Day2 cells had the lowest median arrest coefficient, with very few cells being completely arrested (Figure 3C). These data showed that HSC Day2 cells spent more time at the same location than HSC Day4 and MPP Day2 cells. Interestingly, no cell was continuously motile, and virtually all arrest coefficient values we calculated were higher than 40%, consistent with reports in the literature of HSPCs adopting a ‘stop and go’ motion within the BM parenchyma (Upadhaya et al., 2020).

Because of the high arrest coefficients measured and the observed wiggling motion of many cells, we sought to quantify the linearity of motion of all tracked cells. To do this, we measured two further parameters, confinement index and ratio (Figure 3D-E). The former is the ratio of maximum radius (distance between initial and furthest point reached by a cell) (Shakhar et al., 2005) to the total distance covered by the same cell (path length), and the latter is the ratio of displacement (the distance between the initial and the final positions of each cell) to path length (Supplementary Figure 3). In both cases HSC Day2 cells resulted to be the least linear cells, HSC Day4 cells more variable and linear than HSC Day2 ones, and MPP Day2 cells more linear than HSC Day4 cells (Figure 3D-E). Permutation tests on the two values across all cell groups confirmed heterogeneity in the linearity of movement among different cell types, but no obvious differences between confinement index and confinement ratio for each cell type (not shown). We therefore directly compared the values of maximum radius against displacement for each cell (Figure 3F). As expected, for each cell the maximum radius was equal to or larger than the displacement, and the Pearson correlation coefficient between these two variables was 0.938, which confirmed the strong similarity between confinement index and ratio. Because the confinement ratio would fail to distinguish between cases where a cell moved extensively but eventually returned to its original position and one that moved very little, and because this value is ultimately affected by the length of observation (the longer the observation window, the longer the path), we selected to use only confinement index for further analyses.

Together, our track analysis indicated that more differentiated HSPCs migrated in a faster and more linear way, raising the questions whether (1) there may be intrinsic properties of HSCs and MPPs that may determine the observed behaviour, and (2) it may be possible to deconvolute the differentiation state of HSC Day4 cells. To address the first question, we analysed an existing transcriptomics dataset of HSPCs (Cabezas-Wallscheid et al., 2014), comparing the transcriptome of HSCs against that of combined MPP2/3/4 populations, and we performed a Gene Ontology (GO) analysis of the resulting differentially expressed genes with a focus on GO categories of genes involved in regulating cell migration and adhesion and, as an internal control, proliferation (Sommerkamp et al., 2021). Interestingly, we found genes involved in the first two cellular functions upregulated in both HSCs and MPPs (Figure 3G and H). However, HSCs had a stronger expression of genes involved in cell adhesion and negative regulation of cell migration (Figure 3G), while MPPs presented stronger signatures for positive migration parameters and, as expected based on the biology of these cell populations, proliferation (Figure 3H). Taken together, the results of microscopy and transcriptomic analysis suggest that there is a gradual behavioural evolution from HSCs to MPPs, associated with a differential expression of genes likely to affect the observed cellular behaviours, namely HSCs being slower and less linear, and MPPs being faster and more linear.

### Tracks clustering highlights patterns of HSPC-niche interactions

Because of the observed differential expression of genes involved not only in cell migration but also in cell adhesion, we reasoned that analysis of how tracked cells interacted with their surrounding microenvironment may provide important clues to deconvolve their stem vs. differentiated state. Consistent with previous observations (Khorshed et al., 2015; Lo Celso et al., 2009), HSPCs tracked in endothelial cell and osteoblast reporter mice were found adjacent to vasculature and near osteoblasts, and all tracked cells were near the endosteum (Supplementary Video 3 and Supplementary figure 4A and B). We decided to focus on interactions between HSPCs and nestin-GFP stromal cells because these cells are known components of the HSC niche that have been shown to survive irradiation damage (Asada et al., 2017; Méndez-Ferrer et al., 2010; Pinho et al., 2013) and their spatial organisation is such that it is possible for HSPCs to be in perivascular, endosteal areas but still be localised at a wide range of distances from nestin-GFP stromal cells (Figure 1E and F, Supplementary Figure 1D and Figure 2A-C).

We reasoned that analysis of migration and HSC-niche interactions measured for HSC Day2 and MPP Day2 cells would provide an indication of patterns associated with more and less primitive cells, which could then be useful to interrogate the HSC Day4 cells dataset. As it was apparent from qualitative inspection of the data that very few cells resided in and maintained direct contact with nestin-GFP stroma for long periods of time (Figures 1 and 2, supplementary figure 1D and 2C and supplementary videos 1-4), we hypothesised that residing within a certain radius from nestin-GFP cells may be important instead. To define such radius, we measured the distance between each HSC Day2 and MPP Day2 cell and the closest nestin-GFP stroma over time. MPP Day2 cells showed a trend towards a greater range of distances from nestin-GFP stromal cells, and the average/median distance of both cell groups combined was approximately 25 µm (Figure 4A). When we measured the distance between tracked cells and nestin-GFP stroma over time, MPP Day2 cells were most consistently situated further away from nestin-GFP stroma than HSCs Day2 cells. Indeed, the median distance to nestin-GFP stroma was always greater than 25µm for MPP Day2 cells, while it was sometimes lower and sometimes greater than 25 µm for HSC Day2 cells. In both groups a number of cells remained most of the time within this distance from nestin-GFP stromal cells, while others where consistently beyond, and a smaller proportion of cells moved across this threshold (Figure 4B). Interestingly, no HSC Day2 cells were constantly adjacent to nestin-GFP cells (Figure 4B). Therefore, rather than focussing on direct interactions with these cells, we wondered whether a better measure of HSC Day2 cells interactions with the microenvironment would be their persistence near nestin-GFP cells. We therefore calculated a new parameter to describe the percentage of time during which a cell would reside within 25µm from a nestin-GFP stroma cell, and we called it ‘NestinArrest25’, or ‘NA25’ (Figure 4C). Interestingly, HSC Day2 cells had a stronger bimodal distribution of NA25 values, with most cells showing either very high or very low NA25, while MPP Day2 cells had a higher proportion of cells with intermediate NA25 values (NA25 >20% and <80%) and less than 20% cells with NA25 >80% (Figure 4C). However, a permutation test indicated no statistically significant difference in the NA25 values of HSC Day2 and MPP Day2 cells. Mesenchymal stromal cells secrete a high number of relevant factors, such as CXCL12 and SCF, that can impact haematopoietic cells, including HSCs (Crippa and Bernardo, 2018; Méndez-Ferrer et al., 2010). We therefore wondered if cells that spent most of their time near or further from nestin-GFP stroma cells would present unique migration dynamics. To address this, for each cell we compared their mean speed when they were located within 25 µm from nestin-GFP stroma cells versus when they were further than 25 µm (Supplementary Figure 4C). Cells with NA25=100 are reported along the X axis, and cells with NA25=0 are reported along the Y axis. For all the other cells we observed that the average cell speed was not dependent on their relative distance to nestin-GFP cells, confirmed with a Pearson correlation coefficient of 0.44. Next, to address whether more and less primitive cells would be linked to specific behaviours, we used a UMAP analysis to gain insights on the diversity of the tracks analysed based on a selection of the parameters that described them (Figure 4D). HSC Day2 cell tracks were polarised towards two extremes of the resulting map, with some cells clustering at the top and some at the bottom (Figure 4D, blue circles). MPP Day2 cell tracks had a wider distribution, consistent with the higher variance in the parameters that described them (Figure 4D, mustard dots). Unsupervised clustering analysis grouped all HSC Day2 and MPP Day2 cells into 5 clusters (Figure 4E). Most HSC Day2 cells were found in cluster 3, characterised by lowest NA25, speed, confinement index, standard deviation of speed and, fittingly, highest arrest coefficient, and cluster 4, defined by the highest NA25 and intermediate levels of all other parameters (Figure 4F). On the other hand, MPP Day2 cells were more evenly distributed across all clusters. These data are consistent with the HSC Day2 population being heterogeneous, with cells in different clusters likely being in more and less differentiated states, and raising the question of which cellular behaviour may correspond to stemness.

**Figure 4.**
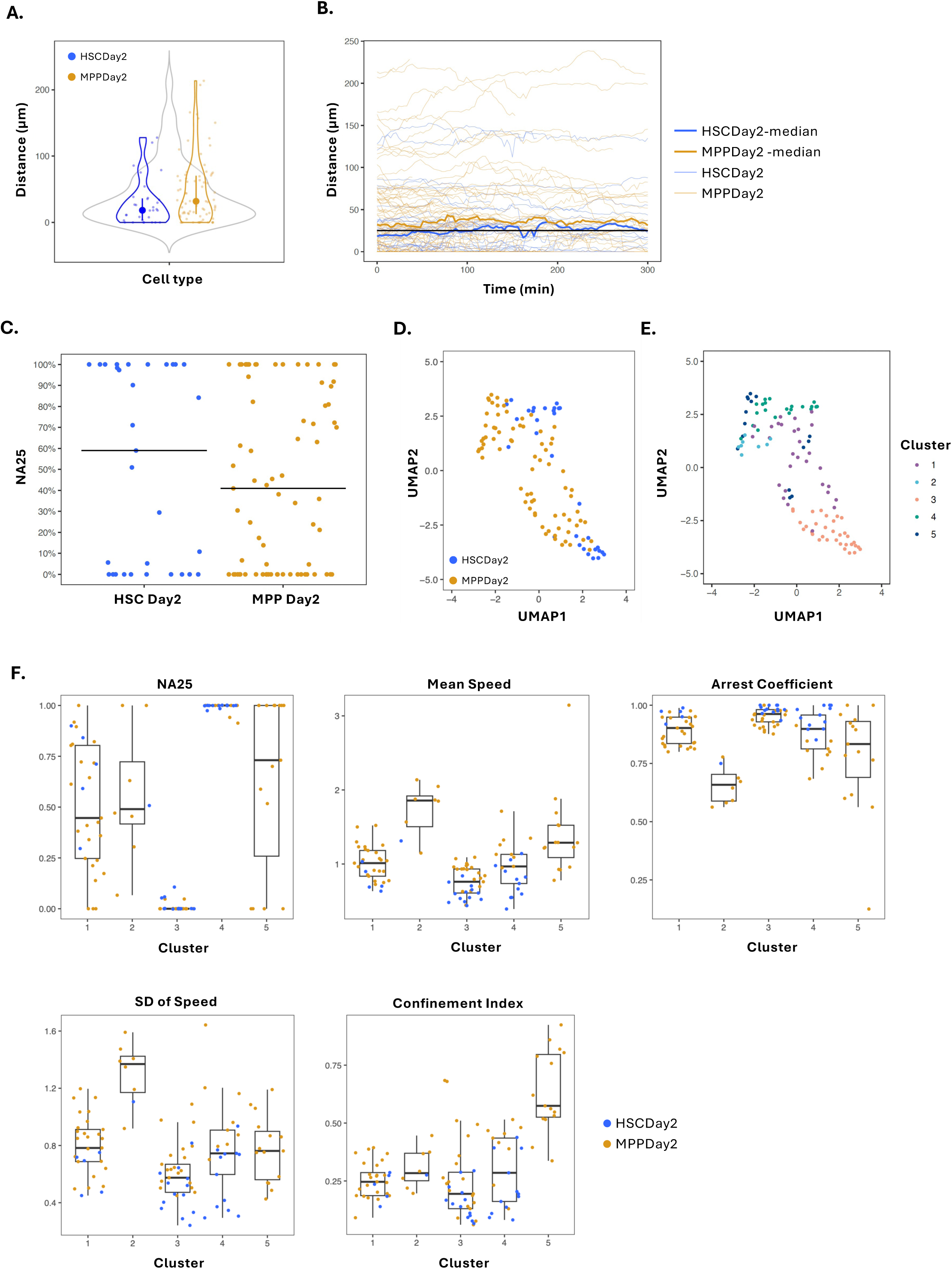
Tracks clustering based on migration and interactions with nestin-GFP stroma cells. **(A)** Distance between each tracked cell to the closest nestin-GFP cell at the start of theirtracks. **(B)** Representation of the distance from each tracked cell to the nearest nestin-GFP cell over the time lapse duration. Thin lines represent each tracked cell, while thick lines represent the median per cell type. The black line highlights 25 µm distance. **(C)** Analysis of nestin arrest 25 coefficient (time spent within 25 µm from nestin-GFP stroma cells, NA25) for HSC Day2 and MPP Day2 cells. Black lines indicates the median value for each population. **(D)** UMAP analysis of HSC Day2 and MPP Day2 cells based on mean speed, standard deviation of speed, arrest coefficient, confinement index, and NA25 coefficient. **(E)** Unsupervised clustering analysis of HSC Day2 and MPP Day2 cell tracks. **(F)** Migration parameters for the cells in each cluster.

### Changes in HSPC-niche interaction patterns correlate with differentiation state

To test whether patterns of interaction with the microenvironment would correspond to differentiation states, we sought to gain insights on the molecular phenotype of a number of HSC Day2 cells that we tracked. We developed a protocol that allowed assessing multiple well-stablished HSC and differentiation markers (CD150, CD48 and a mix of multiple lineage markers) through *ex vivo* whole mount staining of the calvaria of the imaged mice, and measured markers’ expression on the same tomato+ cells that we had imaged (Figure 5A).

**Figure 5.**
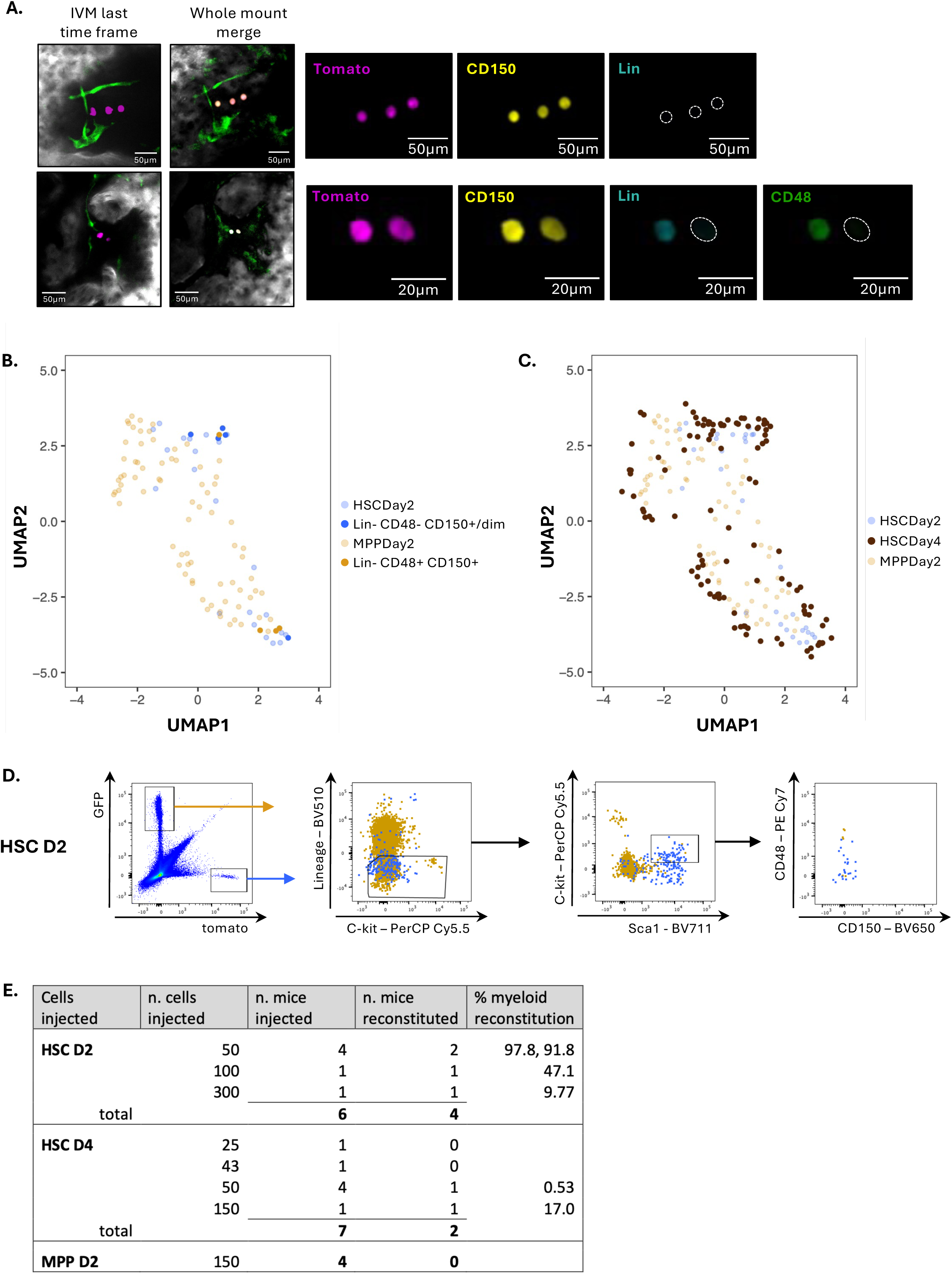
HSPC-niche interaction patterns correspond to differentiation states. **(A)** Two representative images of *ex vivo* whole mount immunostaining of calvarium BM samples from transplanted and imaged mice (HSC Day2). Top and bottom rows include: IVM image (last time frame acquired) and merge whole mount image, followed by higher magnification view of single channels. **(B)** UMAP of all HSC Day2 and MPP Day2 tracked cells, with bright full circles highlighting the cells further analysed by immunostained, colour coded based on resulting primitive (blue) or differentiated (mustard) phenotype. **(C)** Projection of HSC Day4 cells (dark brown circles) onto the cluster map generated from HSC Day2 and MPP Day2 cells (light blue and light brown). **(D)** Representative example of flow cytometry plots and gating strategy showing bone marrow data analysis after IVM of HSC Day2 and MPP Day2 cells. **(E)** Summary table of all transplant results achieved with the three cell populations (HSC Day2, MPP Day2 and MPP Day4).

Consistent with the heterogenous tracks of these cells, we could detect different phenotypes, with some cells appearing more primitive (Lin-CD150+ CD48-), while others already expressed CD48 or markers included in the lineage cocktail, indicating a more differentiated state (Figure 5A). Next, we linked the phenotype of immunostained cells with their tracking data and identified them within the UMAP landscape to infer the relationship between cell behaviour and differentiation state (Figure 5B). All but one lin-CD150+ CD48-HSCs Day2 cells mapped within cluster 4, with the highest NA25. Conversely, all but one of the cells positive for Lin, CD48 or both were within cluster 3, with the lowest Narrest 25 and speed and the highest arrest coefficient.

Finally, to interrogate the diversity of HSC Day4 cells, we mapped their tracks over the existing UMAP of Day2 cells (Figure 5C). Interestingly, HSC Day4 cells did not group into obvious clusters, but rather were scattered across the map. Some cells overlapped with the HSC Day2 cluster 4 cells, others with HSC Day2 cluster 3 cells, others with MPP Day2 cells across the whole map. These results were consistent with HSC Day4 cells being heterogenous, with only a small proportion of cells likely to have remained undifferentiated.

To further validate that HSC Day2 cells included a high proportion of phenotypic HSCs, we performed flow cytometry analysis of bone marrow harvested from calvarium and long bones of some HSC Day2 and MPP Day2 mice (Figure 5D). We observed that all HSC Day2 (tomato+) cells fell within the lineage negative gate, showed various degrees of c-Kit downregulation, as expected post-transplant (Chen et al., 2016; Shen et al., 2004), included the cells with highest c-kit and Sca-1 expression, most did not express CD48, and there were some still present within the SLAM-HSC gate. Instead, MPP Day2 cells (GFP+) were present mainly within the Lineage+ gate, had lower expression of c-Kit and Sca1, and expressed higher level of CD48.

Finally, to validate that HSC Day2 cells contained a substantial proportion of functional HSCs, HSC Day4 a smaller one and MPP Day2 cells did not contain any, we sort-purified tomato+ and GFP+ cells from recipient mice and performed secondary transplantation experiments. We performed blood lineage analysis in peripheral blood every 4 weeks (Supplementary figure 5A) and a final BM analysis at 20-29 weeks after transplant.

Consistent with our microscopy and UMAP analyses, HSCs Day 2 cells led to the highest rate of BM reconstitution, and HSCs Day4 cells to lower ones. MPP Day2 cells were unable to reconstitute any mice in the long term, as expected. Taken together, our data suggest that HSPC-BM microenvironment interactions change as differentiation takes place, and it is possible to infer the degree of stemness of HSC progeny tracked *in vivo* based on the dynamics of their cellular behaviours.

## DISCUSSION

Haematopoietic stem cells are essential to maintain blood production throughout life and a key cell population for therapy of several blood disorders, from leukaemia to genetic and autoimmune diseases. HSC transplantation (HSCT) was the first stem cell-based therapy to be applied in the clinic and remains the only curative option for many patients. Despite its widespread application, HSCT is still highly challenging for patients and unsuitable for older and frailer ones. Understanding the mechanisms underpinning the early stages of HSC-driven haematopoietic regeneration is crucial to accelerate short-term engraftment without hindering long-term HSC function, which would improve HSCT success and increase its applicability. In this study we identify vicinity to Nestin-GFP stroma cells and intermediate values of migration parameters to correlate with primitive HSC phenotypes, and an increasing heterogeneity of cell-microenvironment interactions to underpin HSC progeny differentiation.

The BM is a complex tissue form by different blood, stromal and vasculature cells embedded in ECM matrix (Comazzetto et al., 2021; Pereira et al., 2024). These cells influence HSC numbers, self-renewal and differentiation capacity through both direct cell-cell interactions and the generation of a supportive cytokine milieu (Comazzetto et al., 2021). The coordinated effects of these molecular exchanges are not yet fully understood, partly because still little is known about the temporal nature of the cellular interactions between HSCs and the BM microenvironment. Our finding that HSC progeny resides in the vicinity of nestin-GFP cells, with more primitive HSC progeny spending more time near these cells is consistent with static intravital and *ex vivo* imaging analyses that reported HSCs in close proximity to nestin-GFP periarteriolar stroma in steady state and immediately following transplantation (Itkin et al., 2016; Kunisaki et al., 2013; Méndez-Ferrer et al., 2010). Additionally, it suggests that cytokine production rather than direct cell-cell adhesion may be the main molecular mechanism though which these stroma cells support HSCs and their progeny. Several microscopy studies report HSCs in the vicinity of stroma, vasculature and megakaryocytes (Acar et al., 2015; Kokkaliaris et al., 2020; Lo Celso et al., 2009). The abundance of these cell types seems the main driver of the observed distributions (Kokkaliaris et al., 2020), and it has been argued that in steady state there may not even be cytokine gradients across BM (Kunz and Schroeder, 2019). The relative rarity of nestin-GFP stroma cells and their specific ability to withstand stress such as irradiation conditioning suggests that in these settings they may generate cytokine gradients that in turn affect HSPC’s localisation.

There are several studies that highlighted different molecular regulators of HSC repopulation capacity. Holmfeldt *et al* performed a functional screen on transplanted HSCs and identified genes involved in vesicular trafficking (nbea, cadps2), cell surface receptor turnover (gprasp) and secretion/remodelling of ECM components (crispld1) as important regulators of HSC engraftment (Holmfeldt et al., 2016). These data suggest that HSCs engage in complex, active and bidirectional cross-talk with their niche as they engraft, leading to the hypothesis that HSCs may actively condition the niche (Holmfeldt et al., 2016). These observations are consistent with our finding that HSC reside within more confined niches than MPPs and that the stability of HSC-niche interaction (*i.e.* time spent within the field of view) decreases as differentiation develops. In addition, they also identified Arhgef5, a RhoGEF, is required for HSC repopulating capacity *in vivo* (Holmfeldt et al., 2016). Rho is important for cell rounding especially in amoeboid movement, important in podosome formation. Fitting with this, in our live imaging we observed that all transplanted cells, even the ones that did not migrate during the time of acquisition, were wriggling and generating protrusions to interact with their surroundings.

The migratory nature of HSPCs has been increasingly under scrutiny, and a combination of intravital microscopy reports have been building a picture whereby the most primitive HSCs are least motile, while progenitors are increasingly migratory (Christodoulou et al., 2020; Lo Celso et al., 2009; Secchi et al., 2025; Upadhaya et al., 2020). The fact that in our study the cluster of cells enriched for phenotypic HSCs had slightly higher speed and speed standard deviation and lower arrest coefficient suggests that in transplantation settings the pattern may be different. This would be consistent with our earlier observation that HSCs from mice in the acute phase of *T.spiralis* infection, which showed higher primary repopulation potential, were more migratory and engaged with wider niches within the BM parenchyma of transplant recipients (Rashidi et al., 2014). We recently identified tight interactions between HoxB5-KuO+ vWF-GFP+ HSCs and LepR+ stroma cells, with HSCs preferentially wrapped by LepR+ cell protrusions compared with HoxB5-KuO+ vWF-GFP-HSPCs (Secchi et al., 2025). As LepR+ stroma is severely damaged by multiple stresses, it is likely that this wrapping is reduced in transplant recipients, enabling wider motion. Consistent with this, a recent study were HSCs and MPPs tracked in vitro were reported to be motile, with HSCs showing non-directional movement, while MPPs always moved more linearly (Camargo Ortega et al., 2026). Of note, even though engrafting HSCs may be more motile than the same cells in steady state, MPPs and their progeny continue to show higher motility, and clusters that contain uniquely these cells have the highest mean speed and speed standard deviation and lowest arrest coefficient, or the lowest confinement index.

Transcriptomics analysis of engrafting HSC progeny highlighted a strong program of differentiation within the first week after HSC transplantation. Most homed HSCs became MPPs in this time, with transcriptional changes such as downregulation of self-renewal transcription factors or increase of TF related to myeloid cells and erythro-megakaryocytes observed already 1 day after transplantation (Dong et al., 2020). These data support our observation that already at day 2 and 4 after transplantation we could detect different cell dynamics and interaction patterns, raising the possibility that multilineage differentiation may trigger heterogeneous, likely lineage-specific patterns of HSPC-niche interactions. Of note, the rates of *in vivo* HSPC expansion and *ex vivo* colony forming assay output reported by Dong *et al*. are compatible with the increased number of cells we could detect at 4 vs. 2 days post HSC transplantation and with the decreasing engrafting ability of HSC Day2, HSC Day4 and MPP Day2 cells in transplantation assays. Interestingly, other microscopy-based studies describing haematopoietic engraftment following HSPC transplantation reported the formation of cell clusters that grew in size over time (Lee et al., 2026; Malide et al., 2012). Our work revealed cell clusters of similar size driven by HSCs alone at equivalent time points, allowing a direct assessment of the remarkable functional potential of these cells and highlighting how intravital microscopy of engrafting HSC progeny may be developed into an efficient assay to evaluate the effectiveness of interventions aimed at maximising HSC self-renewal and progeny output in transplantation settings.

## Authors contributions

CLC and KD conceived the project, planned the experimental design and supervised data acquisition and analysis. RK and SGA drove the wet work and data analysis with contributions from GA, BP, AZL, CP and CFL. CL completed all tracking data QC and statistical analyses. MCR and CNW performed bioinformatics analyses. CLC and SGA wrote the first draft of the manuscript and everyone contributed to its editing.

## Acknowledgments

We are grateful to the Imperial CBS and flow cytometry facility and Francis Crick Institute BRF facility for their support of all wet work. This work was funded by Cancer Research UK (Programme Foundation Award C36195/A26770 to CLC with studentship C36195/A27830 to SGA, BBSRC project grant BB_L023776_1, Blood Cancer UK project grant 15040 to CLC and DP, Wellcome Trust Investigator Award 212304/Z/18/Z and 4i clinical PhD studentship to GA, an NIHR Academic Clinical Lectureship to GA, Imperial-ICR Convergence Science Centre project grant P90516 to CP, a BBSRC DTP studentship to BP (supervised by CLC and CFL), by a Royal Society/Irish Royal Society International Exchange Program IEC\R1\180061 to CLC and KRD, and by Science Foundation Ireland Project Grant 18/CRT/6049 to KRD. We are grateful to Dominique Bonnet and Ilaria Malanchi, Francis Crick Institute, and to Paul Bourgine, Lund University, for discussion and support.

## Supplementary Figure legends

**Supplementary Figure 1.**
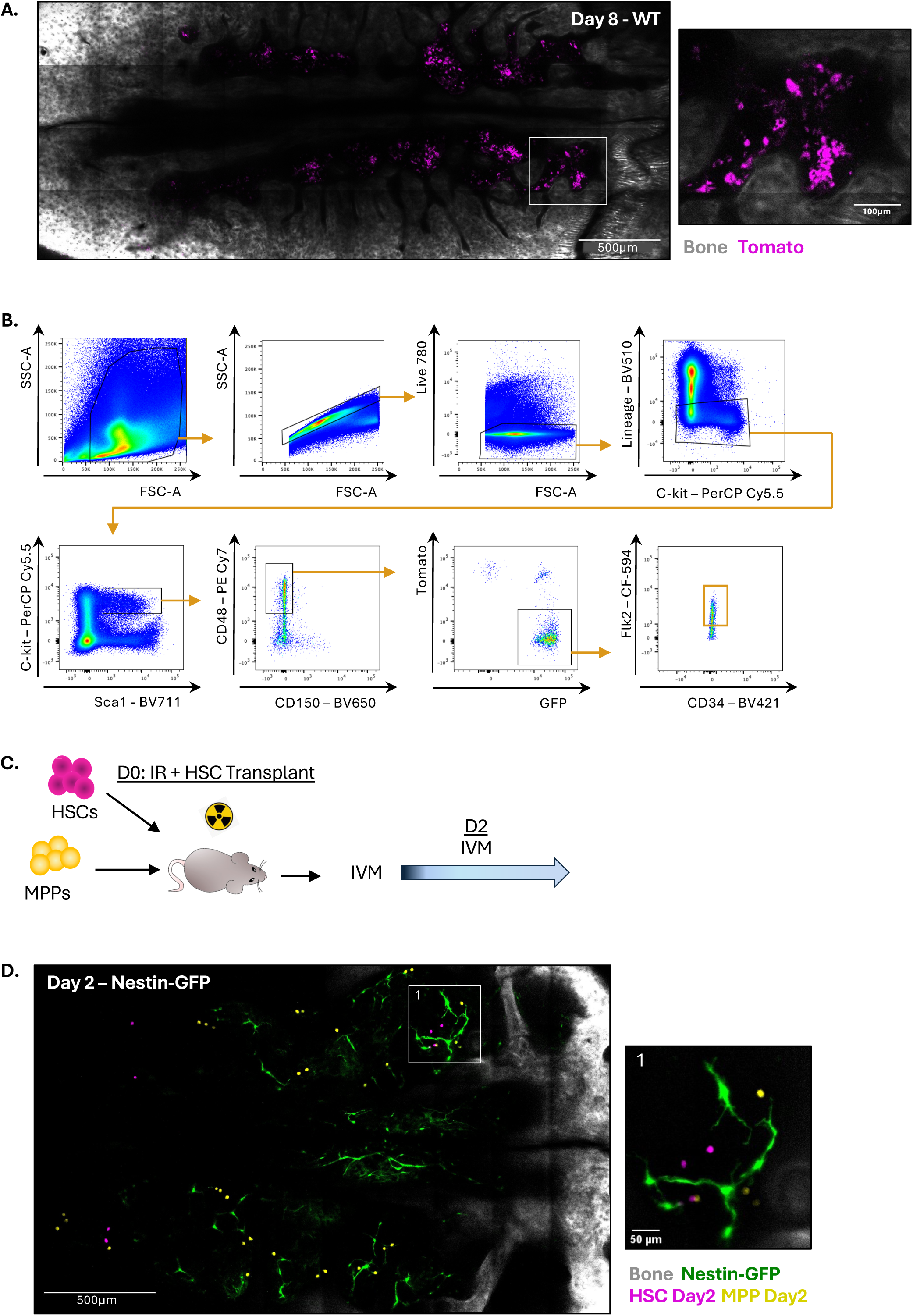
Larger numbers of cells observed by day 8 post HSC injection or day 2 post MPP injection. **(A)** Representative tile scan image of a WT mouse calvarium transplanted with HSCs and imaged at day 8 after transplant. Zoom image on the right side focused on transplanted tomato+ cells. **(B)** Representative example of flow cytometry gating strategy used to sort MPP cells. **(C)** Experimental timeline of sort, transplant and IVM of MPP population. **(D)** Representative tile scan image of calvarium BM of a nestin-GFP mouse transplanted with HSCs and MPPs and imaged at day 2 after transplant. In A and D white framed areas within the tile scan images are shown at higher magnification on the right side to highlight tomato+ cells. White: bone collagen second harmonic generation signal; magenta: tomato. In D green is GFP from nestin-GFP+ stroma and yellow is GFP from MPP progeny.

**Supplementary Figure 2.**
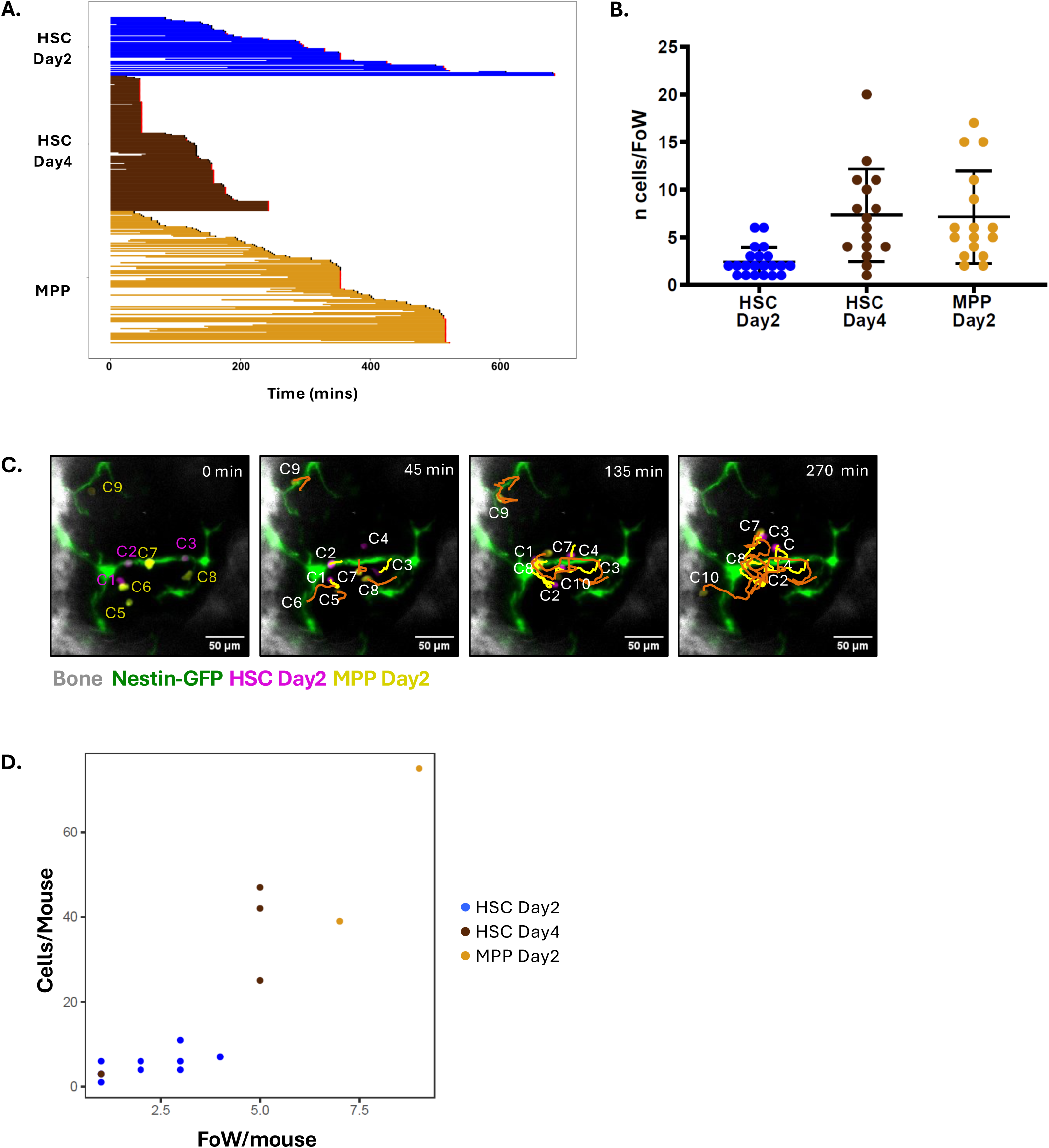
The analysed dataset. **(A)** Swimmer plot of all analysed cell tracks, where each line represents one cell track, and the end of each track is colour coded depending on the imaged cells leaving the field of view before the end of the time lapse (black) or staying in the same field of view for the whole duration of the time lapse (red). Lines starting beyond the zero time point are tracks of cells that appeared in the field of view during the course of the imaging session. **(B)** Number of cells of the different transplanted cell populations per field of view (FoW) acquired by IVM. **(C)** Representative time frames of a time lapse dataset acquired on a nestin-GFP mouse calvarium showing tracks of transplanted tomato+ (magenta) HSC and GFP+ (yellow) MPP progeny at day 2 after transplant. Yellow tracks are of HSC and orange ones of MPP progeny. Grey: bone collagen second harmonic generation signal; green: nestin-GFP signal. **(D)** Dot plot correlating number of cells observed and number of fields of view containing these cells per mouse, per cell type.

**Supplementary Figure 3.**
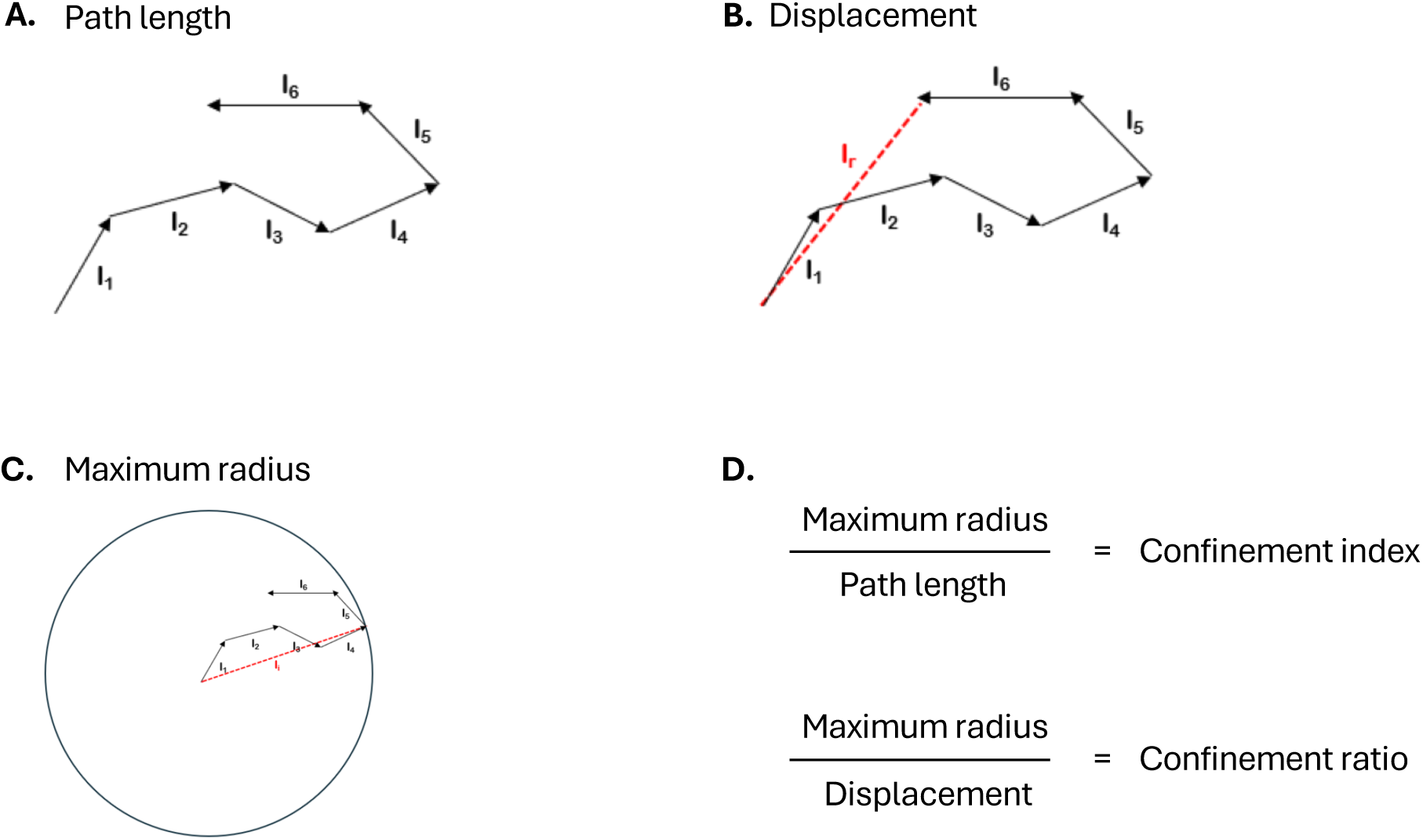
Confinement index and ratio. Schematic representation of path length **(A)**, displacement **(B)** and maximum radius **(C)** measurement for an exemplar track. **(D)** Confinement index and ratio are the ratio of track length over path length and displacement, respectively.

**Supplementary Figure 4.**
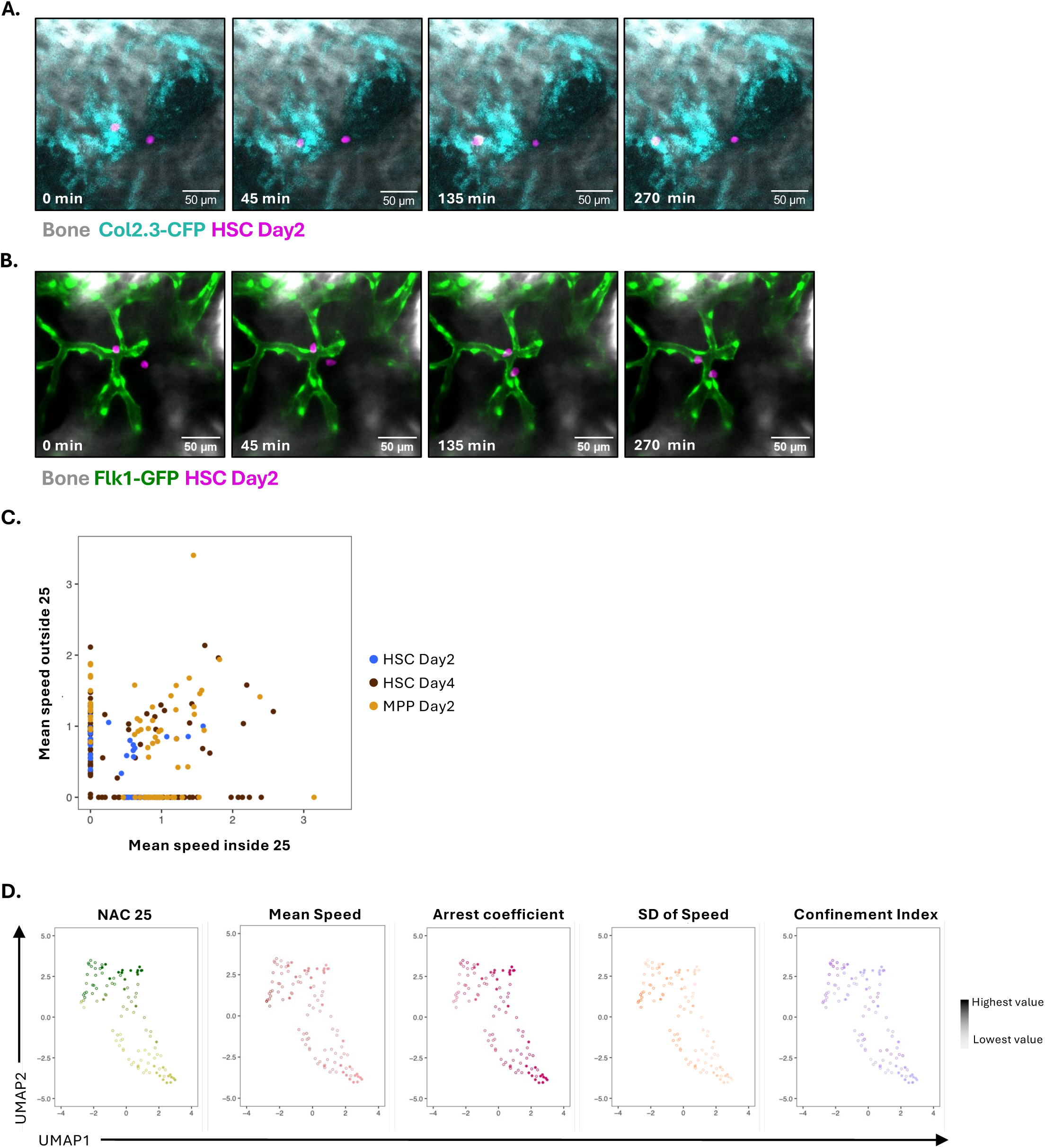
Analysis of HSPC-niche interactions. **(A)** Representative example of HSC Day2 cells observed in the calvarium BM of a Col2.3-CFP reporter mouse calvarium. **(B)** Representative example of HSC Day 2 cells tracked in the calvarium BM of a Flk1-GFP reporter mouse. In (A) and (B) grey is bone collagen second harmonic generation signal, magenta is HSC Day2 signal, cyan CFP, green GFP. **(C)** Mean speed of tracked cells inside and outside a 25 µm radius of nestin-GFP+ cells, colour coded by cell group. **(D)** UMAP representation of all tracked cells, with each cell represented by a dot with colour from dim to bright to represent migration parameter values from low to high.

**Supplementary Figure 5.**
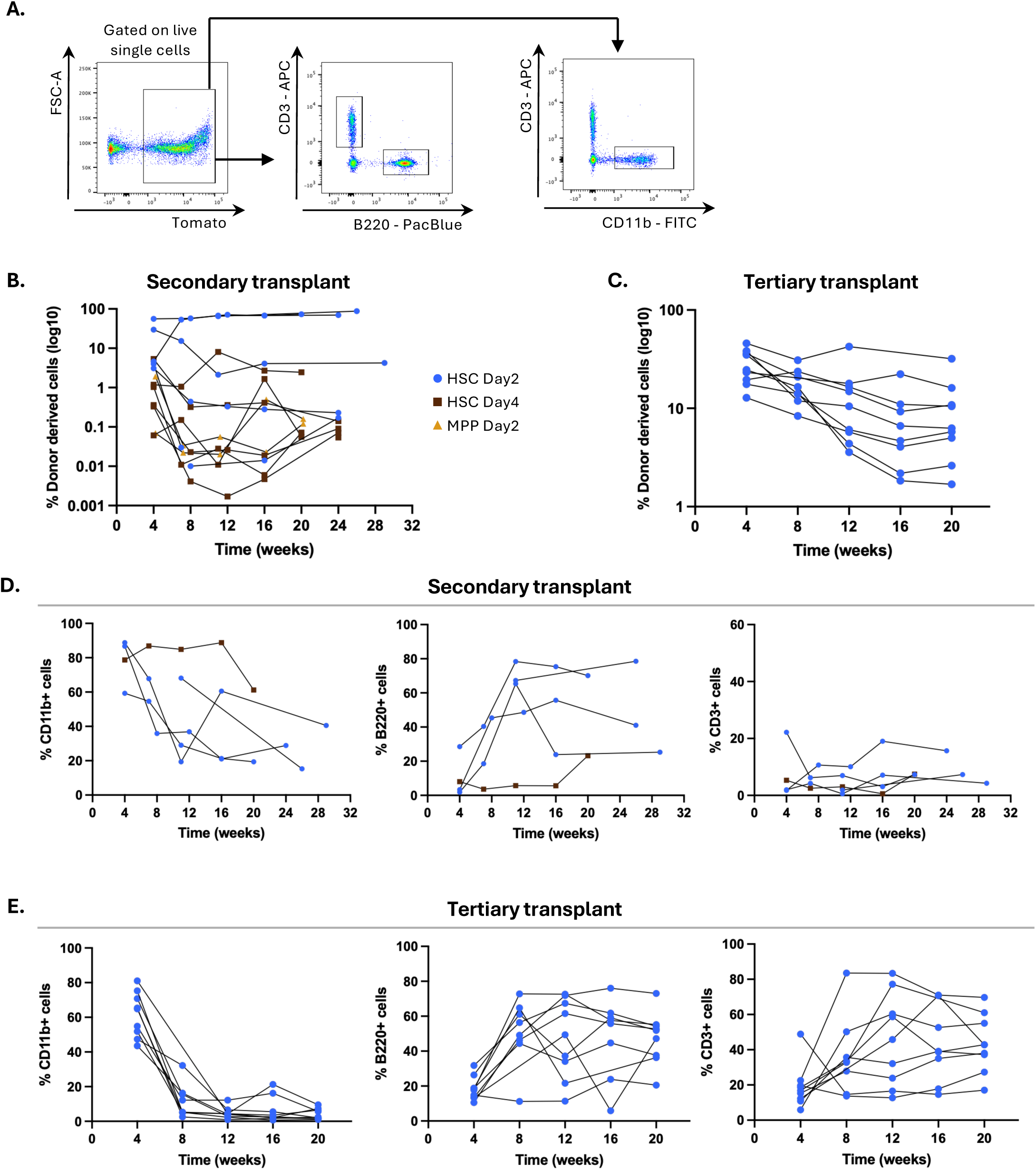
Secondary and tertiary transplants peripheral blood analysis. **(A)** Representative example of flow cytometry plots and gating strategy used to analyse peripheral blood lineage reconstitution in transplant recipient mice. **(B)** Summary plot showing donor-derived haematopoietic reconstitution in secondary recipients. **(C)** Summary plot showing donor-derived haematopoietic reconstitution in tertiary recipients. **(D)** Summary of donor-derived multilineage (T cells, B cells and myeloid cells) reconstitution in secondary recipients. **(E)** Summary of donor-derived multilineage (T cells, B cells and myeloid cells) reconstitution in tertiary recipients.

## Supplementary Video legends

**Supplementary Video 1. HSCs continuously sense the environment through active protrusions.** Time lapse of transplanted tomato+ HSC Day2 cells into a nestin-GFP mouse. Time interval between frames is 3 minutes; total duration of time lapse is 330 minutes. Related to main figure 2A. Red/magenta signal is HSC Day2 cells, green is nestin-GFP, grey is bone collagen second harmonic generation, blue is cy5 dextran, and increasing purple is autofluorescence signal (increasing due to bleaching correction).

**Supplementary Video 2. HSCs present varying degrees of migration capacity through the BM parenchyma.** Time lapse of transplanted tomato+ HSC Day2 cells into a Col2.3-CFP, nestin-GFP mouse. Time interval between frames is 3 minutes; total duration of time lapse is 102 minutes. Related to main figure 2B. Red/magenta signal is HSC Day2 cells, cyan is col2.3-CFP, green is nestin-GFP, grey is bone collagen second harmonic generation, blue is cy5 dextran signal.

**Supplementary Video 3. Transplanted cells intravasate as well as appear or disappear during the time lapse acquisition.** Time lapse of transplanted tomato HSC Day2 cells into a Flk1-GFP mouse (intravasation) and a nestin-GFP mouse (appearance and disappearance from field of view). Time interval between frames is 3 minutes; total duration of time lapse is 252 minutes for the first video and 360 minutes for the second one. White arrowheads indicate the moment a cell intravasates or is not detectable in the field of view anymore. Red/magenta signal is HSC Day2 cells, green is nestin-GFP, grey is bone collagen second harmonic generation, blue is cy5 dextran signal.

**Supplementary Video 4.** Related to main figure 2C. Time lapse of transplanted GFP+ MPP Day2 cells into a nestin-GFP recipient mouse. Time interval between frames is 3 minutes; total duration of time lapse is 516 minutes. Green is MPP Day2 or nestin-GFP, grey is bone collagen second harmonic generation, blue is cy5 dextran signal.

